# Comparative analysis of lifecycle-dependent A to I RNA editing in three members of the *Microbotryum violaceum* fungal complex

**DOI:** 10.1101/2025.01.03.631183

**Authors:** Shikhi Baruri, Alycia Lackey, Joseph P. Ham, Michael H. Perlin

## Abstract

*Microbotryum superbum* (MvSup), *M. intermedium* (MI), and *M. lychnidis-dioicae* (MVLG) are members of the *M. violaceum* fungal complex. Each species infects specific host plant species, resulting in what is commonly known as anther smut. The lifecycle of these basidiomycete fungi includes the haploid, mating, and infection stages. RNA editing is a post-transcriptional process where adenosine (A) is converted to inosine (I) by adenosine deaminase enzymes (ADARs); such modifications to RNAs may lead to synonymous and nonsynonymous codon changes, thereby altering protein function. We observed that 57% to 77% of total editing sites created nonsynonymous codon changes in both haploid and mating stages of the three species.

Moreover, the a2 haploid strain of MI had fewer editing sites compared to other haploid strains. When we compared amino acid substitutions, we found that in both haploids of MvSup and MVLG, Ala was the preferred codon after nonsynonymous codon changes. Among the edited genes, two were edited only at the mating stage in MvSup, undergoing A to I changes within the regions encoding their functional domains. Differential expression analysis revealed that the gene annotated as Apoptosis-inducing factor-1, was upregulated in MvSup at the mating stage, while another gene, for PHB domain-containing protein responsible for cell proliferation, was downregulated compared to the haploid stage. During all stages of the MvSup lifecycle examined, a specific MAPKKK gene was edited in the portion encoding the PKC-like superfamily domain. Also, that gene was edited at a second site during haploid and mating stages but not during the infection stage. Research on RNA editing in basidiomycetes has been limited and is relatively new. RNA editing mechanisms in fungi have been implicated in fungal pathogenesis, although the exact mechanisms and implications remain unclear. Further research is needed to fully understand functional significance of this apparently ubiquitous process in several members of the *Microbotryum* fungal complex, with possible ramifications more generally in fungi.

**Author Summary:** Editing of mRNAs after transcription provides another mechanism for selective expression of, especially proteins, under different stages of development or environmental conditions. Here we report on the characterization of A-to-I RNA editing in three species of the *Microbotryum violaceum* fungal complex, members of the Basidiomycota, where such phenomena have so far been unexplored. We find that such editing is prevalent in different stages throughout the lifecycle of this parasite of plant hosts in the Carnation (Pinks) family. We identified preference for edits that lead to specific amino changes, some of which are limited to one or the other haploid mating-type strains, while others are present preferentially in the mated or plant-infection stages of the lifecycle. Some edits occurred in components of conserved signaling pathways, such as the MAPK pathway, or in genes associated with pathogenicity. Taken together, these results suggest additional hypothesis-driven experiments to further investigate the roles of RNA editing in *Microbotryum*, providing mechanistic insights into the evolution of species in this fungal complex, as well as for those of other pathogenic fungi.

## Introduction

The control of gene expression and the functional diversity of RNA molecules are significantly influenced by the post-transcriptional process known as RNA editing. This process involves altering the sequences of RNA, specifically by converting adenosine (A) to inosine (I) via the action of enzymes known as adenosine deaminases. Regulation of gene expression has been demonstrated to be functionally affected by this mechanism [1]. This kind of RNA alteration can cause nonsynonymous codon changes, that may have an impact on the function of the encoded protein. A-to-I RNA editing has been investigated in mammals for more than 30 years, but it was just recently found in fungi [2]. So far, studies on RNA editing in fungi have demonstrated that A-to-I RNA editing occurs widely in fungi and is not dependent on ADAR enzymes [3]. In Ascomycete fungi like *Neurospora crassa*, A-to-I RNA editing has been shown to be developmentally controlled and adapted for sexual reproduction [4]. Also, there are many selective RNA editing events that have been identified in the medicinal mushroom *Ganoderma lucidum* [5]. Furthermore, it has been suggested in some studies that A to I RNA editing of certain genes may be crucial for the autophagy function during sexual reproduction [6]. A-to-I editing in fungi, such as *Fusarium graminearum*, has been demonstrated to be independent of ADAR enzymes that are typically present in mammals [7]. A significant proportion of pseudogenes are edited to eliminate stop codons, with the majority of the editing process affecting coding areas, which result in amino acid alterations in proteins [7], a common phenomenon in fungi [8]. A-to-I RNA editing has been shown to provide adaptive benefits in fungi, especially in *Fusarium* species, for resolving survival-reproduction trade-offs. The significance of this method in fungal biology is shown by the abundant A-to-I editing sites found in the perithecia of *F. graminearum* [9]. Additionally, experimental validation of the adaptive benefit and functional importance for A-to-I RNA editing in fungi highlights the role this mechanism plays in fungal gene regulation and evolution.

A group of over fifteen basidiomycete fungal species known as the *Microbotryum violaceum* complex infects a similarly large group of plant host species. Each species in this complex shows significant post-zygotic isolation which is evidence of great genetic differentiation and specialization [10, 11]. The complex has been found to infect over 100 species in the Caryophyllaceae family and based on patterns of host specificity and morphological variation, it is considered to be a species complex [10, 12].

The research on A-to-I RNA editing in fungi has revealed its widespread occurrence and developmental regulation, particularly in the context of sexual reproduction. Here, we focus on three members of the *Microbotryum violaceum* complex: *Microbotryum superbum* (MvSup), *M. intermedium* (MI), and *M. lychnidis-dioicae* (MVLG). These three species provide an excellent opportunity to understand the molecular processes regarding sexual reproduction, host preference, and adaptability. Understanding how A-to-I RNA editing happens in these fungi could potentially elucidate the genetic and molecular basis of reproductive isolation and speciation in this complex [13, 14].

The members of the *Microbotryum violaceum* complex have different stages of their life cycle, including the haploid, mating, and infection stages. The infection process starts when diploid teliospores germinate on an appropriate plant surface. Following meiosis, these spores create linear tetrads of haploid basidiospores that reproduce via budding to yield yeast-like sporidial cells. Such cells can mate with partners of opposite mating type. After developing conjugation bridges, mated cells differentiate into dikaryotic hyphae that can penetrate the host tissue after receiving appropriate cues from the host plant. The infection becomes systemic and the hyphae proceed to the floral primordia. After this stage nuclear fusion takes place and the hyphae develop into diploid teliospores. Finally, these teliospores replace the pollen in the anthers of the flowers. The cycle is repeated through transmission by pollinators who spread the spores to uninfected flowers [15].

In this paper we examined A to I RNA editing for MvSup, MI, and MVLG in three different stages of their lifecycles to understand how A to I RNA editing varies among these members of the fungal species complex. Our research aims to identify the commonalities as well as distinctive variations in RNA editing within this particular fungal group. Additionally, in Basidiomycota, RNA editing may be required to grow in a different environment or substrate, a feature possibly associated with fungal pathogenesis, although the effect of protein functions after editing is still unclear [16]. Thus, our research should lead to better understanding of how the editing process influences the expression of pathogenicity related genes in these fungi. With this information, strategies could be developed to prevent or lessen the effects of fungal diseases on hosts, which is crucial for managing ecosystems and agriculture.

### Overall Analysis pipeline

In this paper editing sites of RNA were identified and compared across different species. In the three species, their comparison was performed in two stages – two haploid stages with different mating types and a mating stage. In two species, the infection stages were also compared. Within species, all stages were compared to identify commonly edited genes. The genes RNA-edited in just one of its stages but not in the other stages were considered stage-specific or uniquely edited genes (Fig 1).

**Fig 1.**
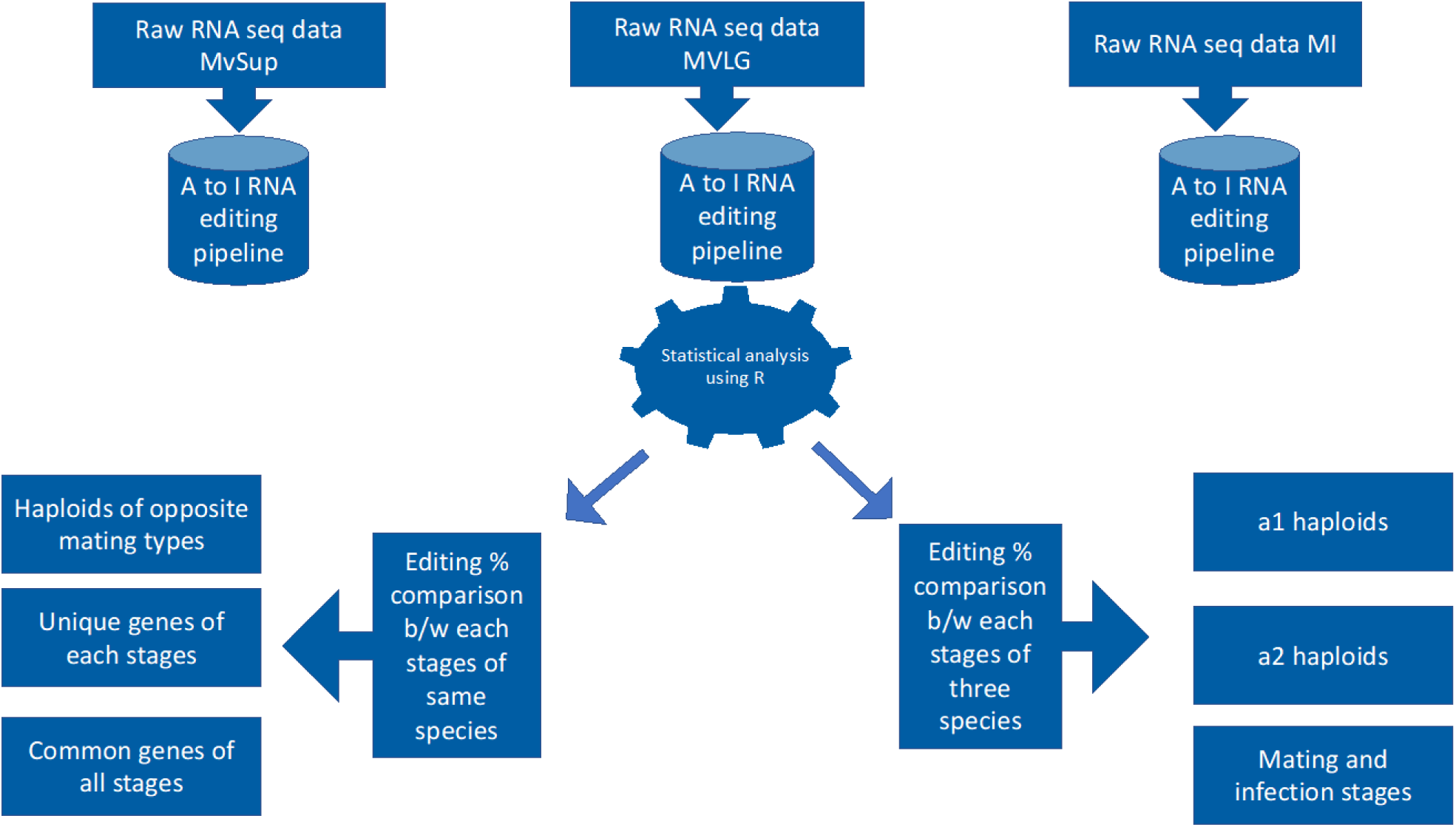
The overall approach to compare RNA editing sites across the species and stages of the *Microbotryum* life cycle. We illustrate the data used from three species and the two types of comparisons made within and between species.

## Results

### A low number of A-to-I RNA editing sites were detected in MI

We have developed a pipeline to identify and analyze the RNA editing sites in mRNA of MvSup, MI and MVLG, each requiring its own specific host species to complete its lifecycle. The data were analyzed for three different stages of the lifecycle of these fungi. MVLG p1A2 haploid strains had more editing sites than MVLG p1A1 all other haploid strains of two species MvSup and MI. Also, MI haploid strains had fewer edited sites than all other haploid strains of MvSup and MVLG. In fact, among all three species of *Microbotryum*, the a2 haploid stain of MI displayed the lowest number of editing sites (Fig 2A).

**Fig 2.**
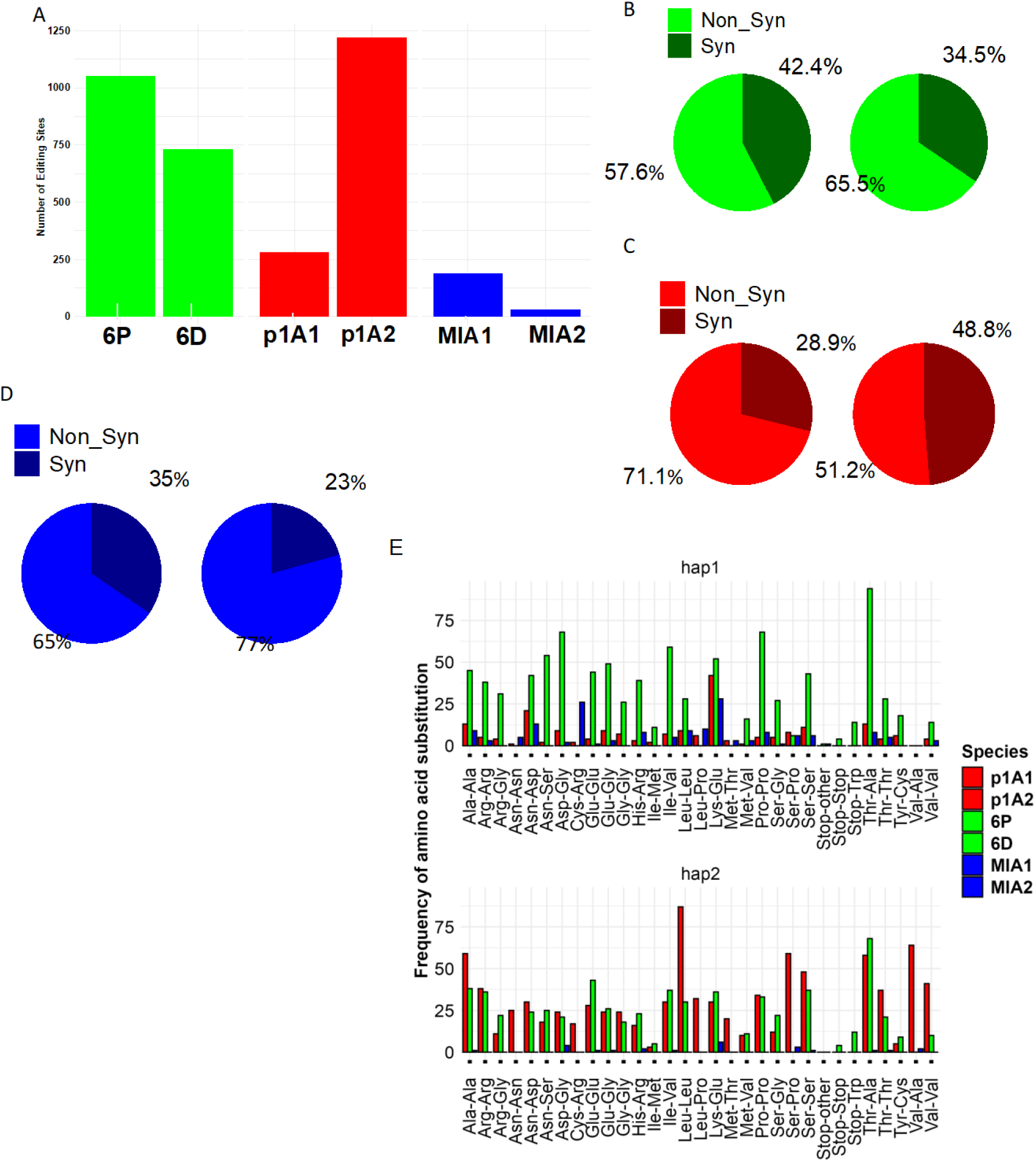
**A)** Comparison of number of editing sites in haploid strains of three species. **(B)-(D)** represent synonymous vs nonsynonymous codon changes: Panel B) represents MvSup haploids 6P (left) vs 6D (right).

Within each species the proportion of A to I RNA edits differed between haploid types. In MvSup, 6P had a significantly higher proportion of edits than 6D (6P proportion of edits = 0.562, CI: 0.547-0.577). In MVLG, the p1A1 had a significantly lower proportion of edits than p1A2 (0.291, CI:0.280-0.302). For MI, MIA1 had a significantly higher proportion of edits than MIA2 (0.792, CI:0.751-0.828). Additionally, all three species differed from each other in the proportion of edits in haploid types (all z-ratios > 8.812, all p-values < 0.0001) (Supplementary S1 Table). The differences in the proportion of A-to-I RNA editing among opposite mating types in *Microbotryum* may reflect some important factors, including genetic drift and natural selection that influenced the evolutionary history of the *Microbotryum* complex. The directionality and extent of these differences in A-to-I editing between these haploid types may differ between species depending on their distinct evolutionary trajectories and environmental setup. This difference may be due to variations in their hosts, which can significantly influence pathogenicity. For instance, one species might infect a broad range of generalized plant hosts, adapting to diverse environmental conditions. In contrast, other species might specialize on a specific host, thriving only in a narrow ecological niche.

### Three types of amino acid substitution are prevalent in the haploids of *Microbotryum spp*

Nonsynonymous changes were more frequent in all species and both haploid types (all differences in non-synonymous and synonymous changes were significantly different from 0), except in MVLG p1A2 which had equal numbers of nonsynonymous and synonymous changes (estimated mean difference in nonsynonymous and synonymous changes: -0.0492, CI overlapped 0: -0.161 - 0.0631). In the MvSup strain 6P, 363 and 688 genes were synonymously and nonsynonymously edited, respectively, and in 6D, 310 and 421 genes were synonymously and nonsynonymously edited, respectively (Fig 2B). In MVLG p1A1, 82 and 202 genes were found synonymously and nonsynonymously edited, respectively, while in p1A2, 595 and 625 genes were found synonymously and nonsynonymously edited, respectively (Fig 2C). In MIA1, 65 and 124 genes were synonymously and nonsynonymously edited, respectively, and in MIA2, 7 and 23 genes were synonymously and nonsynonymously edited, respectively (Fig 2D).

When we compared the predicted amino acid substitutions of each haploid strain of all three species, we found mainly three types of substitutions common to all three a1 mating-type strains (p1A1, 6P, and MIA1): threonine (Thr) to alanine (Ala), asparagine (Asn) to aspartic acid (Asp) and lysine (Lys) to glutamic acid (Glu). .

Interestingly, Thr to Ala amino acid substitutions were also prevalent in the a2 mating-type haploid strains of MvSup and MVLG. In addition, cysteine (Cys) to Arginine (Arg) and Lys to Glu were prevalent in MIA1 and A2, respectively. Valine (Val) to Ala conversion was the most frequent amino acid substitution in the p1A2 (a2 mating type) strain. Serine (Ser) to Proline (Pro) conversion was also prevalent in the p1A2 strain of MVLG. Moreover, synonymous and nonsynonymous codon changes occurred in STOP codons of both mating types of MvSup. We observed STOP to Trp codon conversion in 6P and 6D. However, only one event of STOP to Gly conversion was observed.

Synonymous codon changes also occurred in both mating types in codons for Ala, Thr, Pro, Glu, Val, and Ser (Figs 2E). An analysis of the preferred codons for MVLG revealed that for the most degenerate position (i.e., third position), seventeen out of eighteen had a G or C base [15]. Therefore, synonymous codon changes primarily influence the GC composition of the edited genes. That suggests RNA editing is essential for preferred codon selection based on GC composition in *Microbotryum spp*.

Green color represents nonsynonymous codon changes and dark green color represents synonymous codon changes. Panel (**C**) represents MVLG strains p1A1(left) vs p1A2 (right). Red color represents nonsynonymous codon changes and maroon color represents synonymous codon changes. Panel (**D**) represents MIA1(left) vs MIA2 (right). Blue color represents nonsynonymous codon changes and dark blue color represents synonymous codon changes. Panel **(E**) represents frequency of different types of codon changes in two mating types of all three species. In this figure hap1 represents a1 mating types and hap2 represents a2 mating types.

### Analysis of unique RNA editing sites in haploids and mating stage of life cycle

Among all the RNA editing sites we identified for MvSup, 243 and 45 unique genes were edited nonsynonymously in a1 and a2 mating types, respectively. Likewise, for MVLG, 161 genes were edited nonsynonymously that are unique for the a1 mating type and 126 unique genes were edited nonsynonymously in the a2 mating type. Similarly, in MI, 43 and 22 unique genes, respectively, were edited nonsynonymously in the a1 and a2 mating types. When we compared the amino acid substitutions, we found that in both haploids of MvSup and MVLG, Ala was the shared preferred amino acid substitution after nonsynonymous codon changes for both species (Figs 3A, B and 4A, B). However, in the a1 mating type of MI and both haploids of MvSup, Arg was another preferred codon for nonsynonymous codon changes. In M1A2, Glu and Gly were the preferred codons for nonsynonymous codon changes (Figs 3C and 4C).

**Fig 3.**
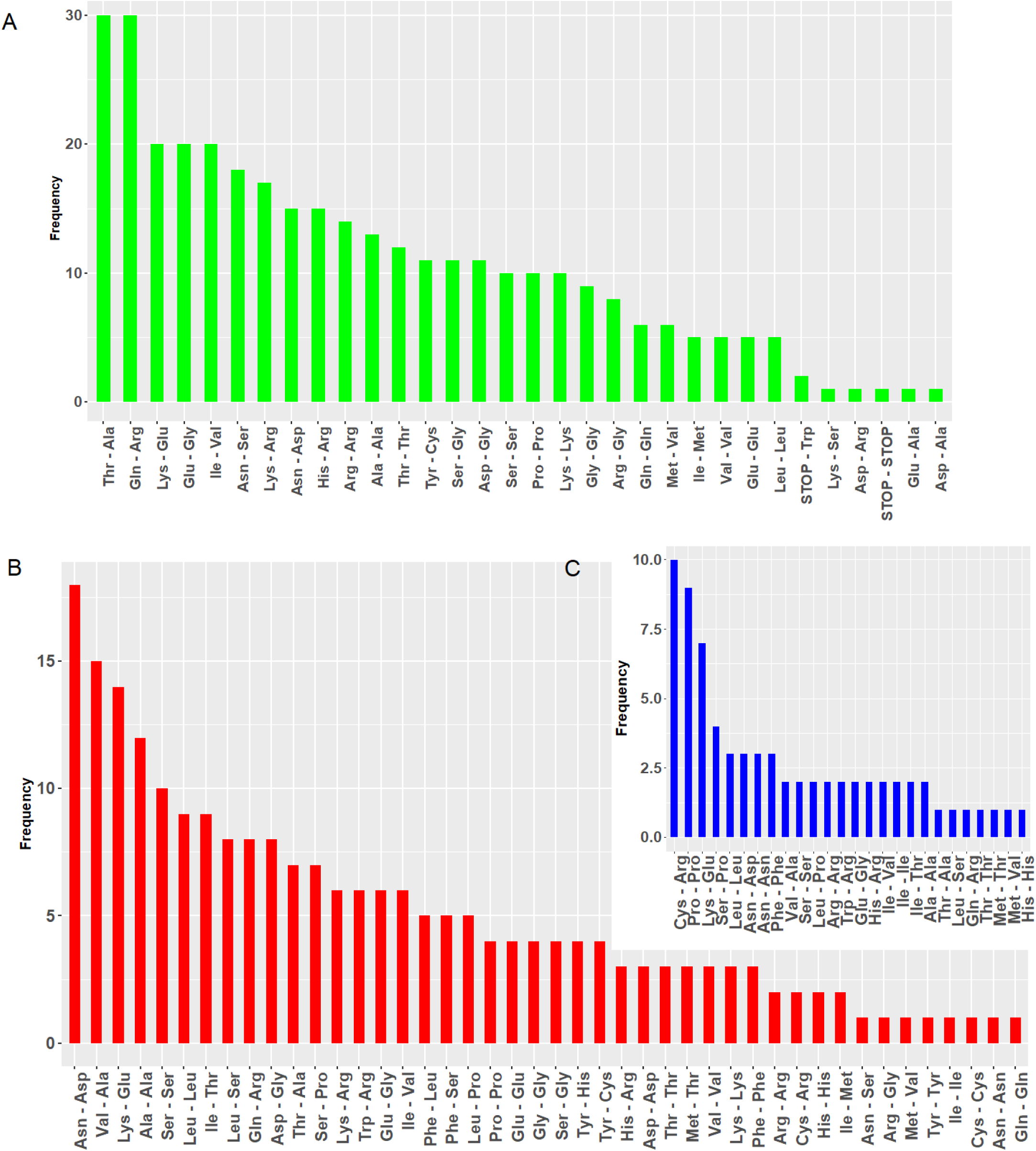
Amino acid substitutions in the unique genes of a1 haploids of three species. A) represents codon conversion of MvSup-6P (green) B) represents codon conversion of MVLG-p1A1 (red) and C) represents codon conversion of MIA1 (blue).

**Fig 4.**
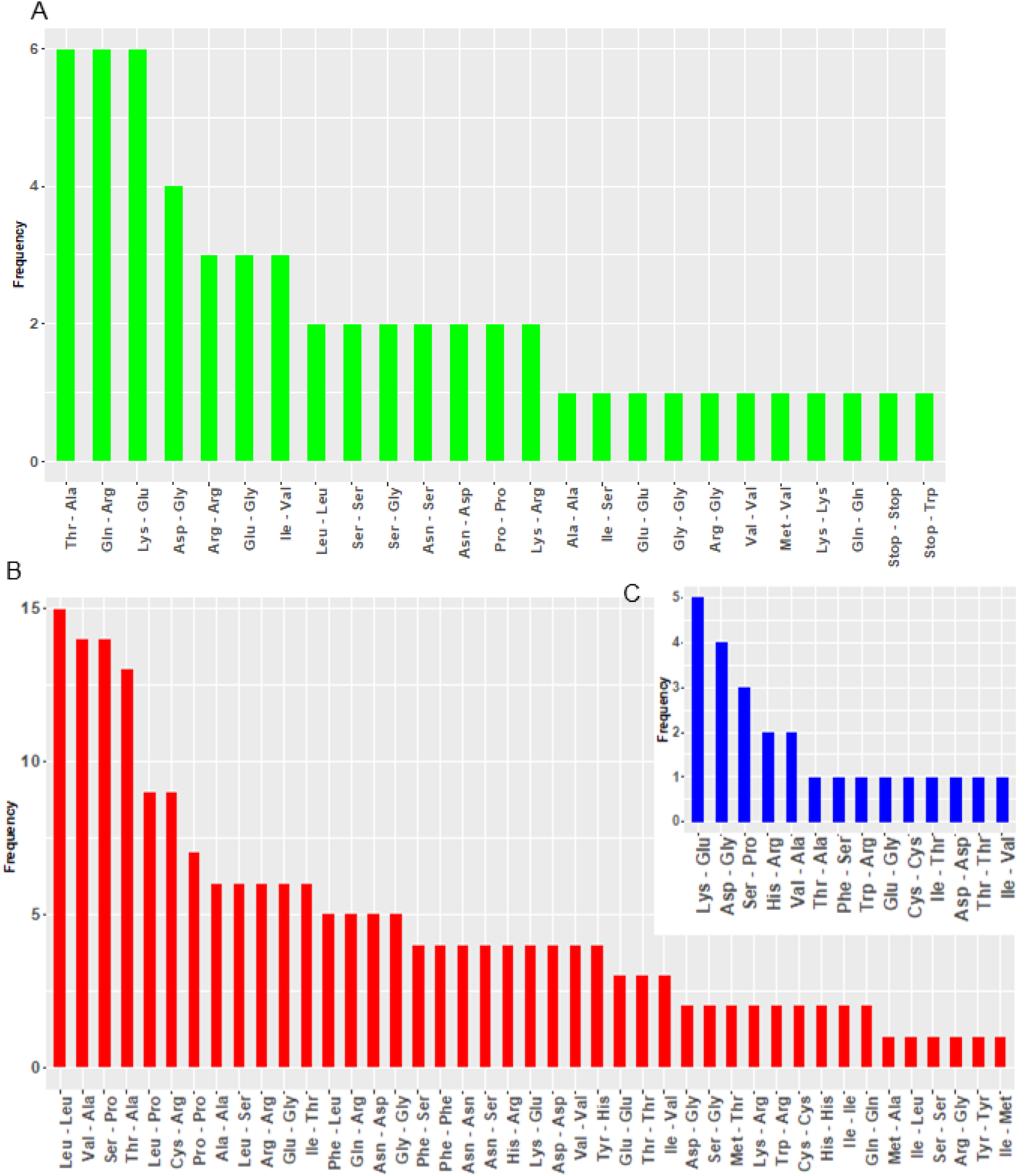
Amino acid substitutions in the unique genes of a2 haploids of three species. A) represents codon conversion of MvSup-6D (green); B) represents codon conversion of MVLG-p1A2 (red); and C) represents codon conversion of M1A2 (blue).

### Important genes edited uniquely at the mating stage of MvSup

In the mating stage of MvSup, we found that a nonsynonymous codon change occurred in a thioredoxin-like protein at codon position 18, which leads to a change from Thr to Ala within the functional domain of the protein (Figs 4A). Also, nonsynonymous codon changes occurred only at the mating stage in the PHB domain-containing protein (g10725) that caused Asn to Ser and Met to Val changes, respectively. Our gene expression data revealed that genes upregulated in the haploid stage tend to be downregulated in the mating stage and vice versa (Fib 4B). For instance, the PHB domain-containing protein was downregulated during the mating stage compared to haploid stages. Another gene (g6828) that codes for a protein that is involved in Phyto steroid metabolic process also was upregulated only during the mating stage. A gene (g3537) that codes for the protein involved in regulation of apoptotic process was upregulated during the mating stage when compared with the haploid conditions (Fig 5B). Also, A to I RNA editing occurred in a gene (g8181) that codes for pheromone-dependent signal transduction involved in conjugation with cellular fusion (Supplementary S2 Table).

**Fig 5.**
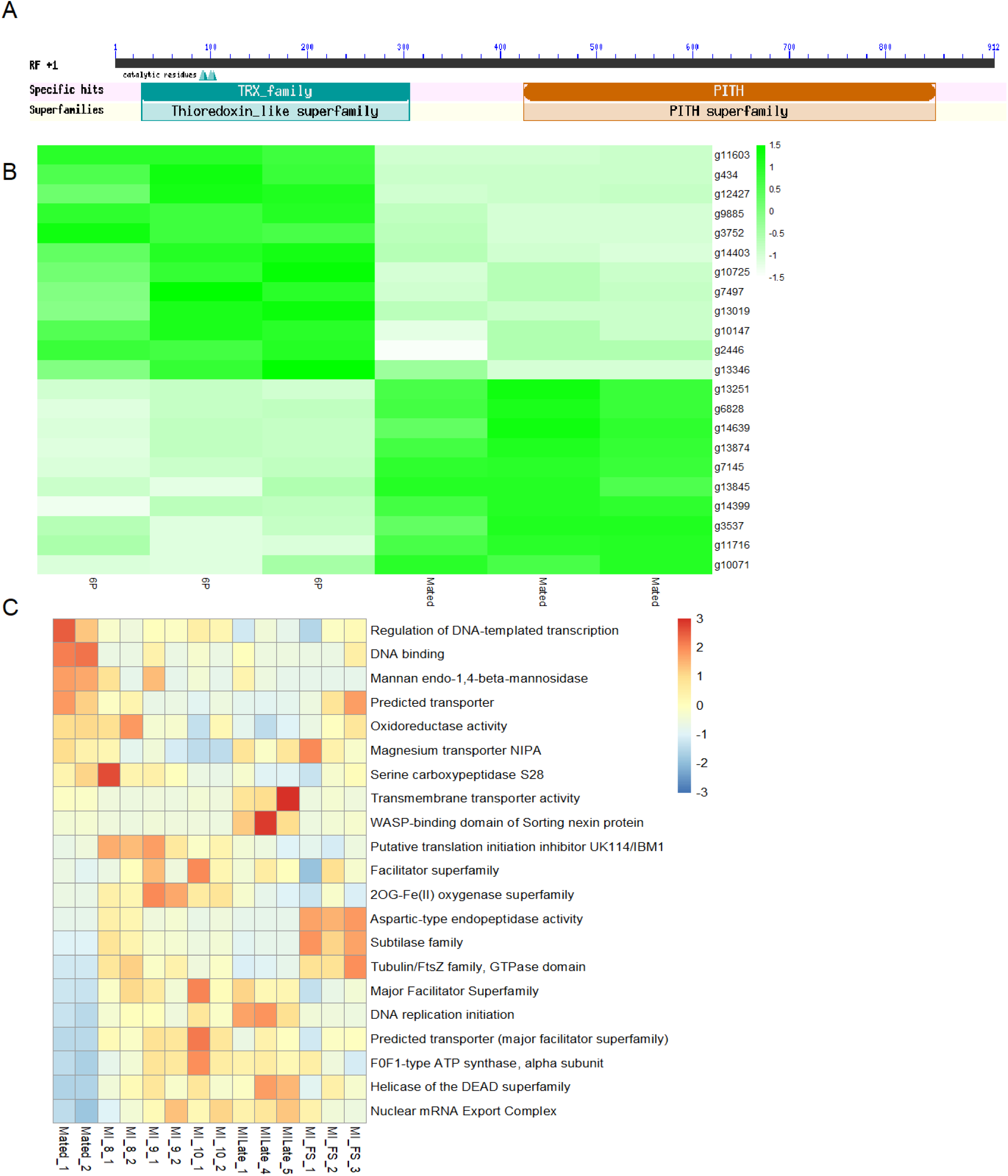
**A**) Thioredoxin-like protein according to NCBI CDS domain identifier. **B**) Represents unique genes for MvSup at the mated stage that were either up or downregulated in the haploid stage. Shade of green indicates degree of up- or down-regulation; **C**) Unique genes for MVLG at the mated stage that were either up or downregulated in the infection stage, based on color scale inset. x-axis represents samples collected at the mated stage (first two samples from left) and at different stages of infection (last 12 samples on the right).

### RNA editing of Subtilisin-like proteases was found only at the mating stage of MVLG, but not in other stages of the lifecycle

Thirty-two genes which are unique for MVLG at the mated stage were either up- or downregulated in the infection stage (Figs 5C, Table 1). For example, one gene associated with serine-type endopeptidase /Subtilase activity was downregulated in the mating stage. RNA editing of that gene at the mating stage caused codon changes from Thr to Ala at position 681. The role of subtilisins in plant pathogenic fungi has been highlighted, suggesting their important role in plant infection [17, 18]. Also, we observed that genes related to transport were downregulated during the mating stage when compared with the infection stage. Moreover, genes associated with DNA binding and those involved with regulation of DNA-templated transcription activity were strongly upregulated in the mated stage and generally had no change in expression in the infection stage.

**Table 1.** Genes uniquely edited in MVLG at mated stage.

| Gene ID | Functions | Codon changes |
| --- | --- | --- |
| MVLG_02763 | Subtilase family | Thr681Ala |
| MVLG_05629 | Predicted transporter (major facilitator superfamily) | Tyr12Tyr |
| MVLG_04456 | UNKNOWN | Val79Ala |
| MVLG_05664 | aspartic-type endopeptidase activity | Ile221Val |
| MVLG_03646 | UNKNOWN | Ser484Pro |
| MVLG_05707 | oxidoreductase activity | Ala508Ala |
| MVLG_03240 | Tubulin/FtsZ family, GTPase domain | Ser31Ser |
| MVLG_02490 | The nuclear mRNA Export Complex | Gln6Gln |
| MVLG_06505 | F0F1-type ATP synthase, alpha subunit | Gln74Gln |
| MVLG_01859 | Predicted transporter (major facilitator superfamily) | Ile186Thr |
| MVLG_06406 | 2OG-Fe(II) oxygenase superfamily | Val256Ala |
| MVLG_04367 | Serine carboxypeptidase S28 | Asn144Asp |
| MVLG_04451 | UNKNOWN | Ile35Thr |
| MVLG_05343 | Predicted transporter ADD1 (major facilitator superfamily) | Ser26Pro |
| MVLG_03076 | Helicase of the DEAD superfamily | Val227Ala |
| MVLG_05391 | Magnesium transporter NIPA | Ser1194Pro |
| MVLG_05353 | UNKNOWN | Asp18Asp |
| MVLG_07033 | UNKNOWN | Leu140Pro |
| MVLG_07149 | DNA binding | Ser334Pro |
| MVLG_02891 | Putative translation initiation inhibitor UK114/IBM1 | Lys91Arg |
| MVLG_04688 | UNKNOWN | Trp87Arg |
| MVLG_00763 | Major Facilitator Superfamily | Leu288Pro |
| MVLG_07198 | DNA replication initiation | Ser579Pro |
| MVLG_06531 | UNKNOWN | Ile349Ile |
| MVLG_00922 | Predicted transporter (major facilitator superfamily) | Gly206Gly |
| MVLG_01981 | UNKNOWN | Ser324Pro |
| MVLG_02652 | WASP-binding domain of Sorting nexin protein | Gln649Gln |
| MVLG_02745 | UNKNOWN | Ser118Pro |
| MVLG_03470 | UNKNOWN | Trp125Arg |
| **MVLG_06263** | DNA binding | Leu100Ser |
| **MVLG_06542** | mannan endo-1,4-beta-mannosidase | Trp136Arg |
| **MVLG_06801** | UNKNOWN | Ala190Ala |

### Common genes edited in haploids and mating stages of MI

We observed very few editing sites in common for the genes of M1A1, M1A2, and mated stages (Fig 6). Of these 7 common genes, some had only nonsynonymous changes across all stages and some of had only synonymous changes across all stages. For example, nonsynonymous editing was observed with the genes associated with gag-polyprotein putative aspartyl protease and synonymous changes were observed with the genes associated with protein phosphorylation activity in all stages of MI. Only 2 genes had different types of changes across stages. The gene coding for Endonuclease-reverse transcriptase was edited nonsynonymously in the mated stage, while codons changed synonymously in the other two stages (Fig 6A and 6B).

**Fig 6.**
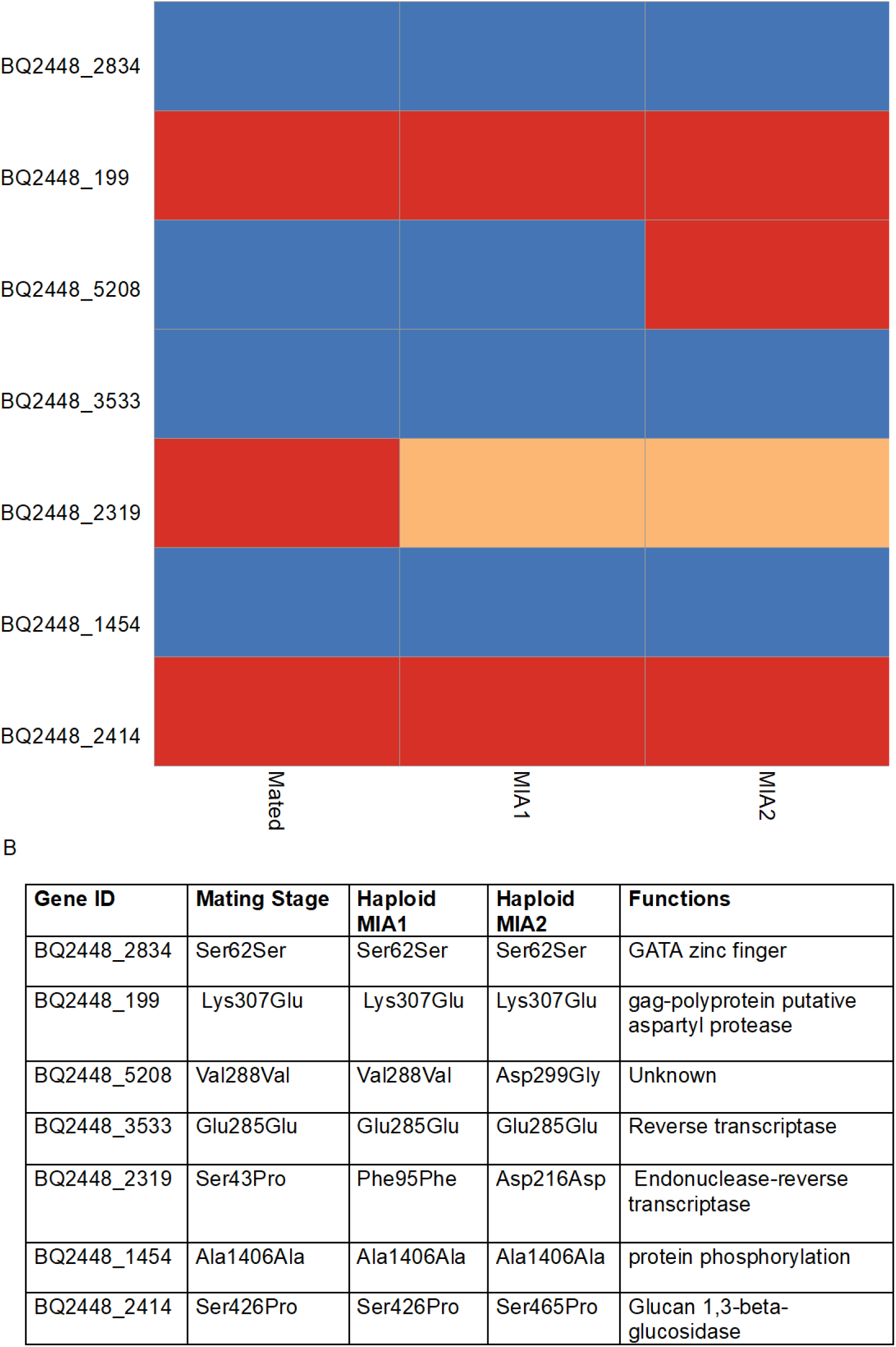
A) Synonymous vs. nonsynonymous codon changes occurred in the genes edited in M1A1, M1A2, and the mated strains of MI. Blue represents synonymous codon changes. Red represents nonsynonymous codon changes. Orange represents synonymous codon changes but different amino acid position. B) This table represents functions of all the genes that are commonly edited in two haploid stages and mating stage of MI.

### Unique genes edited in haploids and mating stages of MI

In these analyses, we identified genes that were edited only at the mating stage. Interestingly, two proteins, according to Gene Ontology (GO) annotations, play a pivotal role in vesicle-mediated transport (GO:0016192) and aspartic-type endopeptidase activity (GO:0004190) (Table 2). It has been suggested that endosomal and extracellular vesicle-mediated transport mechanisms are involved in the secretion of small RNAs at the fungal-plant interface (Kwon et al., 2019). We found that genes coding for Obsolete oxidation-reduction process were uniquely edited in M1A1 and M2A2. This process is a fundamental biological process that plays a crucial role in physiological responses to environmental stimuli. Notably, the genes that are uniquely edited in different stages and caused nonsynonymous codon changes have a variety of biological functions. These genes code for proteins involved in proteolysis and cell redox homeostasis. These functions are essential for maintaining cellular integrity and response to environmental stresses. Functions such as the activity of oxidoreductase and cysteine-type peptidase play a significant role in proteolytic pathways and redox reactions, which are crucial for cellular metabolism as well as defense from oxidative damage.

**Table 2.** GO Annotation of the edited genes unique in haploids and mating stage of MI.

| GO Annotation (M1A1) | GO Annotation (M1A2) | GO Annotation (Mating stage) |
| --- | --- | --- |
| GO:0006508 Proteolysis |  | GO:0006508 Proteolysis |
| GO:0006457 Protein folding |  | GO:0006457 Protein folding |
| GO:0016491 Oxidoreductase activity | GO:0016491 Oxidoreductase activity | GO:0016491 Oxidoreductase activity |
| GO:0008270 Zinc ion binding | GO:0008270 Zinc ion binding | GO:0008270 Zinc ion binding |
| GO:0015074 DNA integration |  | GO:0015074 DNA integration |
| GO:0008234 Cysteine-type peptidase activity |  | GO:0004190 Aspartic-type endopeptidase activity |
| GO:0003676 Nucleic acid binding | GO:0003676 Nucleic acid binding | GO:0003676 Nucleic acid binding |
| GO:0055114 Obsolete oxidation-reduction process | GO:0055114 Obsolete oxidation-reduction process |  |
| GO:0008152 Metabolic process | GO:0008152 Metabolic process | GO:0008152 Metabolic process |
|  |  | GO:0016192 Vesicle-mediated transport |
| GO:0016272 Prefoldin complex |  |  |
| GO:0006633 Fatty acid biosynthetic process |  |  |
|  | GO:0035023 Regulation of Rho protein signal transduction |  |
|  | GO:0005085 Guanyl-nucleotide exchange factor activity |  |
|  | GO:0005515 Protein binding | GO:0005515 Protein binding |
|  |  | GO:0003968 RNA-dependent RNA polymerase activity |
|  |  | GO:0051082 Unfolded protein binding |
|  |  | GO:0055085 Transmembrane transport |
|  | GO:0005509 Calcium ion binding |  |
|  | GO:0035556 Intracellular signal transduction |  |
|  | GO:0045454 Cell redox homeostasis |  |
| GO:0003824 Catalytic activity | GO:0003824 Catalytic activity |  |
| GO:0005506 Iron ion binding |  |  |
|  | GO:0003779 Actin binding |  |
|  |  | GO:0005524 ATP binding |

### Analysis of shared gene functions across life stages in MvSup and MVLG

Where RNA editing was found in all stages of the fungal lifecycle of MvSup and MVLG, genes associated with transport were commonly observed as being edited (Figs 7A and 7B). Distribution of amino acid substitutions in all the common genes edited during haploid, mating, and infection stages of MvSup and MVLG can provide insights into the evolutionary history of *Microbotryum*. We can infer evolutionary relationships, adaptation processes, and the conservation of specific amino acids in critical protein regions by comparing the substitutions between these species. We observed that within the common genes, for those edited in all stages of MvSup, Thr to Ala substitution was the most preferred, while Lys to Glu was preferred for MVLG (Figs 8C and 8D).

**Fig 7.**
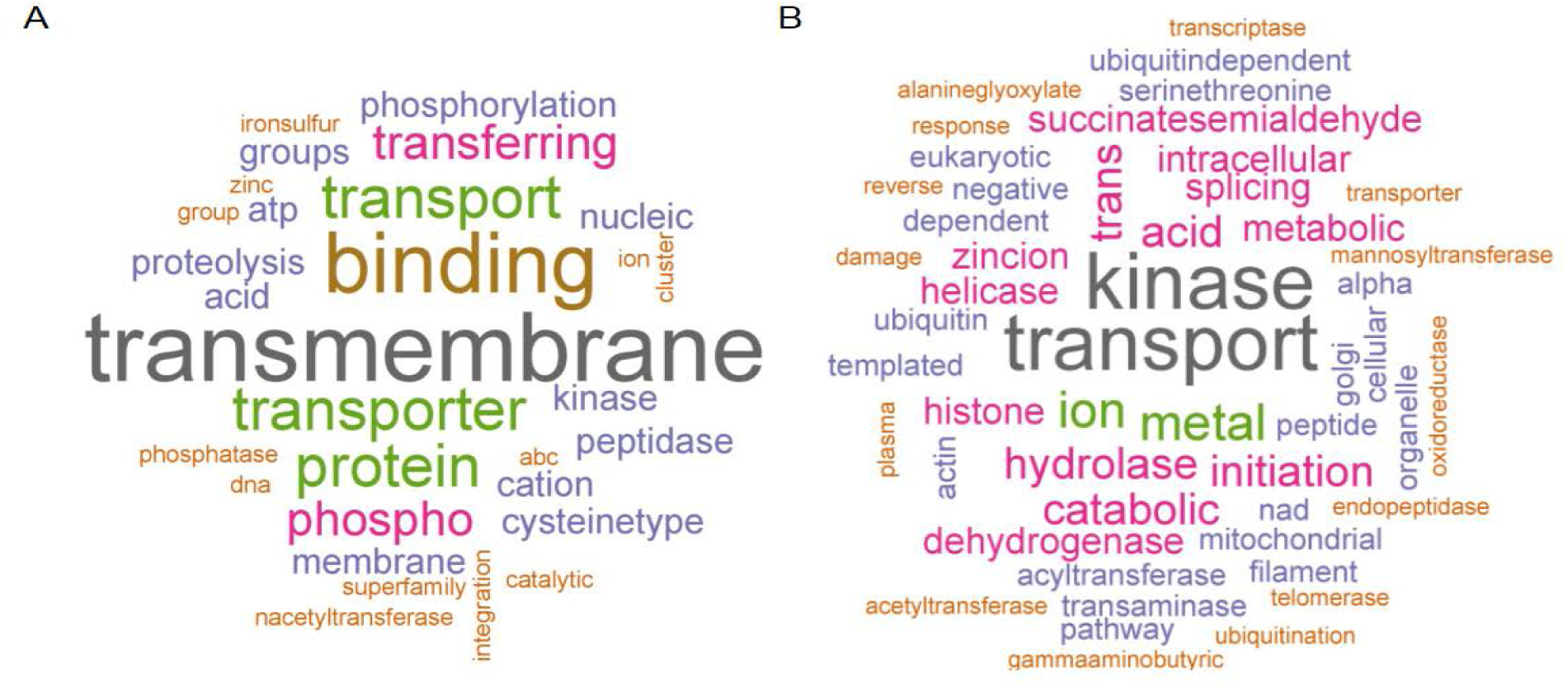
**A)** Word cloud plot of the function of the genes that were edited during haploids, mating, and infection stages of MVLG. **B)** Word cloud plot of the function of the genes that were edited during haploids, mating, and infection stages of MvSup.

**Fig 8.**
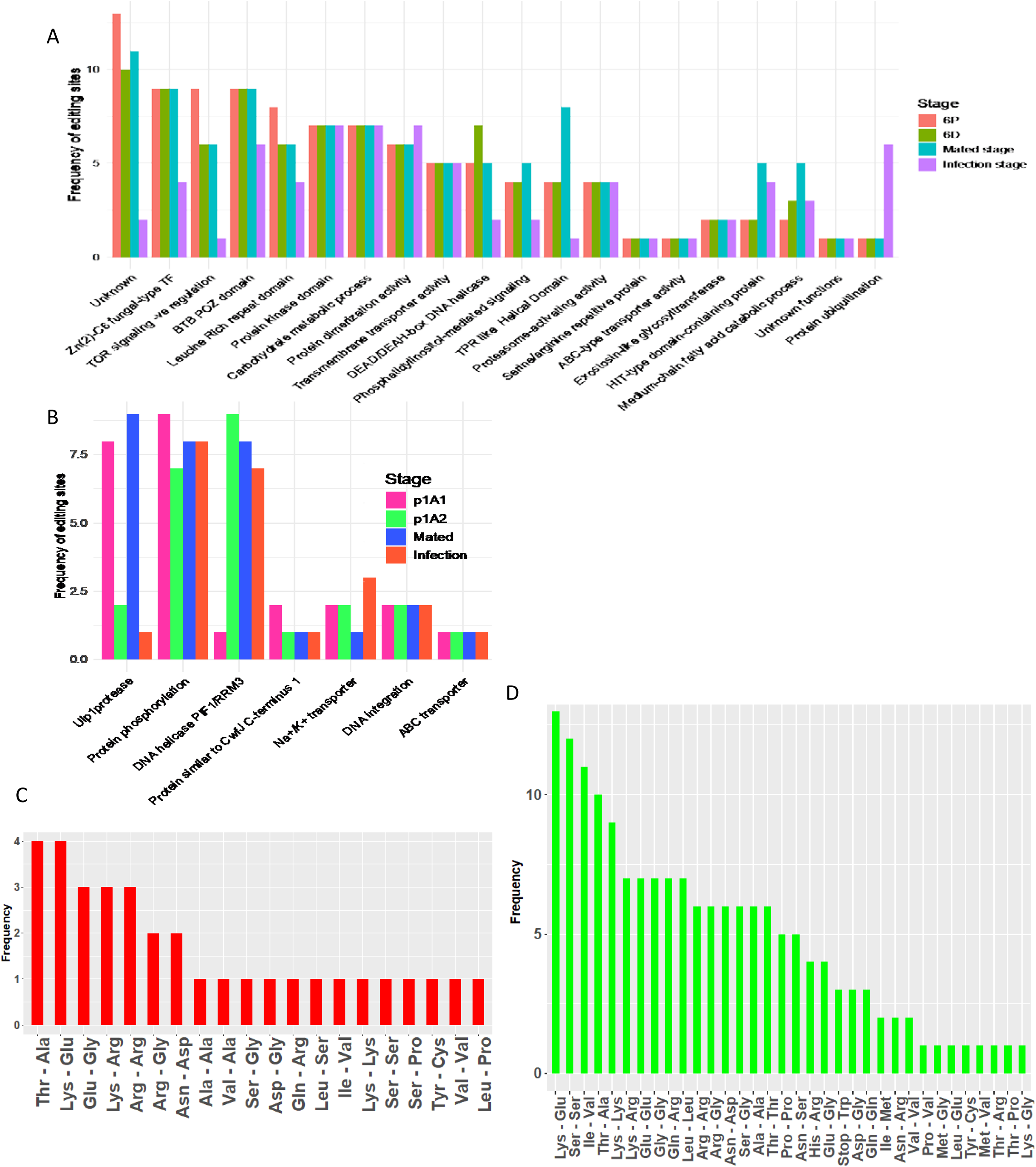
**A)** and **B)** Frequency of editing sites were compared between the common genes that are edited in all three stages (haploid, mating and infection stages) of the lifecycle of MvSup and MVLG, respectively. **C)** and **D)** Represents the distribution of amino substitutions in genes that are common for all three stages (haploids, mating and infection stages) of MVLG and MvSup, respectively.

We identified that among all the genes where RNA editing occurred in every stage of the fungal life cycle in MvSup, five genes had more editing sites than those found for other genes edited in common for all stages of lifecycle in MvSup. These five genes code for BTB POZ domain containing protein, Leucine Rich repeat domain, protein kinase domain, carbohydrate metabolic process, and Zn (2)-C6 fungal-type transcription factor. Among all the common genes, the genes involved in protein ubiquitination and protein dimerization activity had more editing sites during the infection stage compared to all other stages. Also, three genes had more editing sites in the mated stage compared to all other stages; these genes code for the protein associated with phosphatidylinositol-mediated signaling, the protein containing a TPR-like Helical Domain, and a protein containing a HIT-type domain (Fig 8A). In both haploid mating-types, as well as during mating and infection stages of the MvSup lifecycle, a specific MAPKKK gene was edited in the portion encoding the PKC-like superfamily domain. Also, that gene was edited in a second site during the haploid but not in the mating and infection stages (Supplementary S3 Table).

In MVLG, a gene that encodes a Ulp1 protease, a member of the ubiquitin-like protease 1 (Ulp1) cysteine protease family, had more editing sites in the mated stage compared with other stages. This protein plays a crucial role in fungal mating and is responsible for the maturation of SUMO and the processing of the precursor of the ubiquitin-like Smt3 protein into its mature form, as well as the cleavage of Smt3 from its conjugates [19, 20]. Moreover, a gene that codes for Na+/K+ transporter protein had the most editing sites during the infection stage compared to other stages in MVLG (Fig 8B).

### Transport-Related Genes: A Shared Editing Profile Between MvSup and MVLG

Genes associated with transport were the most commonly edited across all lifecycle stages examined for both MVLG and MvSup. Of particular note, a gene that codes for an ABC type transporter was edited in all stages in both species (Figs 8A and 8B). ABC-type transporters are a crucial component of drug resistance and detoxification mechanisms in fungi. These transporters, which belong to the ATP-binding cassette (ABC) superfamily, are involved in efflux-mediated resistance to antifungal agents [21, 22] .

### Comparison of late stage male infected flowers (MILate) vs late stage female-infected flowers (Filate) in MVLG

There were 566 editing sites in MIlate of MVLG and 302 editing sites in FIlate. Among the 107 shared genes, 29 and 30 genes were unique in the MIlate stage and FIlate stage, respectively (Supplementary S4 and S5 Tables). A gene encoding lytic transglycolase was edited only in the FIlate stage. The lytic transglycolase plays a crucial role in the infection process of fungi in plants [23]. The synthesis of these lytic enzymes is usually influenced by environmental stimuli, which leads to fungi changing the composition of their cell walls and their breakdown processes. This makes it possible for the fungus to enter the plant and get the nutrients it needs to continue growing inside the host [24, 25]. We also observed in the MIlate stage, a unique edited gene for methylenomycin: H+ antiporter, a protein found to play a significant role in the salt tolerance and infection process of fungi in plants [26–28] (Supplementary S6 Table).

An additional question was whether genes undergoing RNA editing were differentially expressed at the late infection stage in male flowers compared to the haploid stage of MVLG. We observed among the two genes associated with the Major facilitator superfamily (MVLG_05383 and MVLG_01436), MVLG_05383 was upregulated in the late infection stage in male flowers and the other was upregulated in haploids. The genes coding for flavin-binding monooxygenase, and for CDC24, which has a PB1 domain, were upregulated in the haploid stage compared to the infection stage (Supplementary S1Fig; Supplementary S7 Table). Furthermore, the MIlate stage-specific genes where RNA editing sites were identified encoded helicase, endopeptidase and Major Facilitators (Fig 9A).

**Fig 9.**
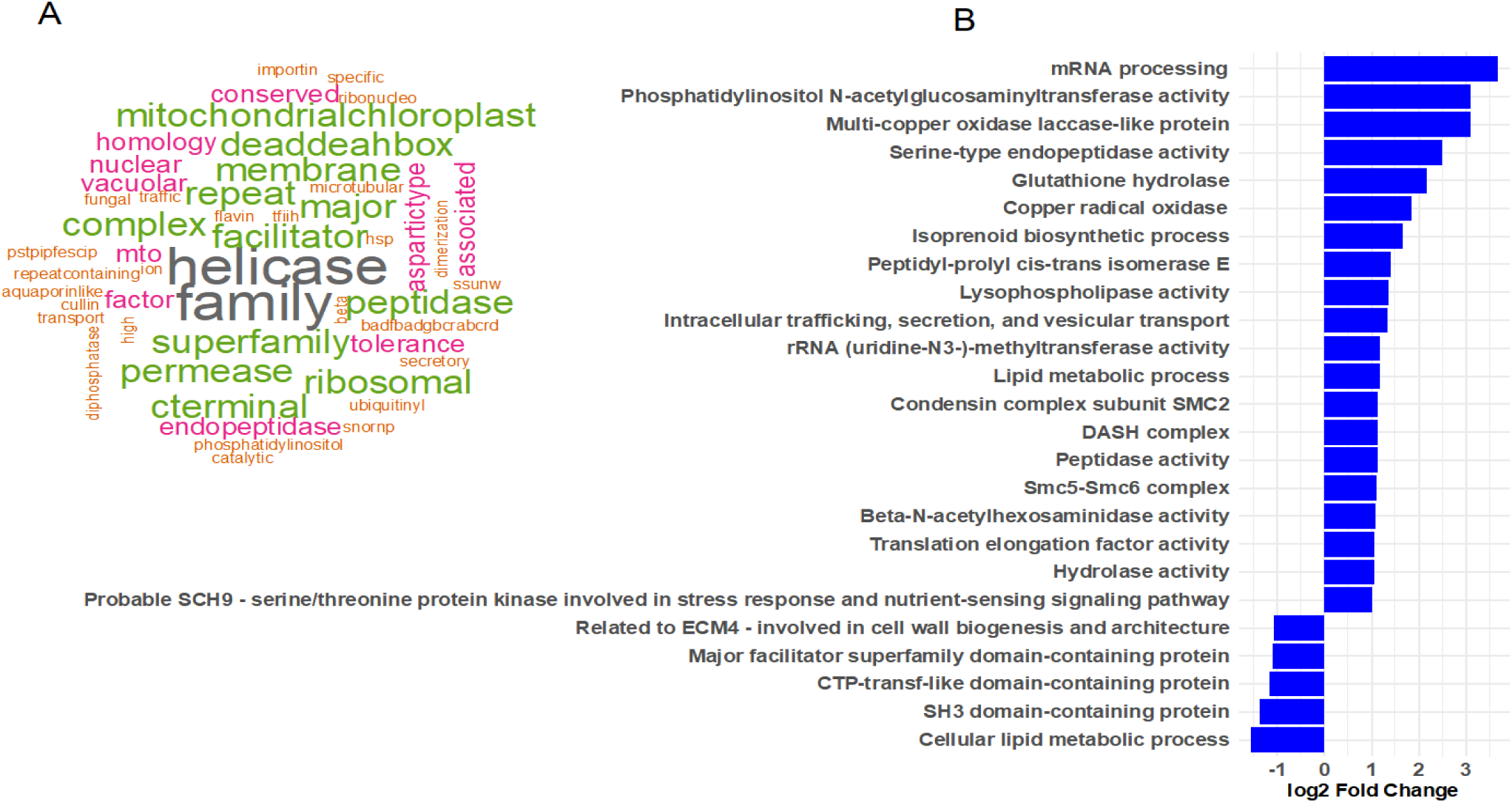
**A)** Word cloud plot of the function of the genes edited only during MIlate infection stage in MVLG. **B)** Differential expression level (mated vs infection stage) of the genes uniquely edited during the infection stage.

### Uniquely edited genes during infection stage of MvSup

Differential gene expression between mated and infection conditions was examined for RNA edited genes of MvSup that occurred only during the infection stage (Fig 9). We identified upregulated genes associated with functions like Isoprenoid biosynthetic process, Probable SCH9 - serine/threonine protein kinase involved in stress response and nutrient-sensing signaling pathway, and Intracellular trafficking, secretion, and vesicular transport. At the same time, ECM4 like protein, which likely contains subtilisin-like serine proteases and SH3 domain-containing protein, were downregulated during infection conditions (Fig 9B). Among all the genes edited only at the infection stage of MvSup, a few genes were identified: a gene coding for PB1 domain containing protein; a gene associated with the regulation of response to osmotic stress; and, a gene involved in phytochelatin metabolism to counteract the host plant’s defense mechanisms, including the production of phytochelatins (Supplemental material). In fungi, proteins containing the PB1 domain are involved in defense responses against different stressors [29], as well those associated with fungal pathogenesis [30]. The PB1 domain is vital for regulating plant responses to stressors and diseases [31]. The domain participates in forming protein complexes for cellular signaling pathways [32].

## Discussion

We performed A-to-I RNA editing analysis, using our in-house pipeline (Fig 10), which includes several functions from the GATK workflow. No specific tool is available for A-to-I RNA editing, and most of the tools that exist are developed for human or mouse models. For validation of the editing sites identified by our pipeline, we used the REDI tool to verify the comparison for one of the species, MI, which showed around eighty percent similarity to the REDI tool results when used to verify the RNA editing sites (Supplementary material).

**Fig 10:**
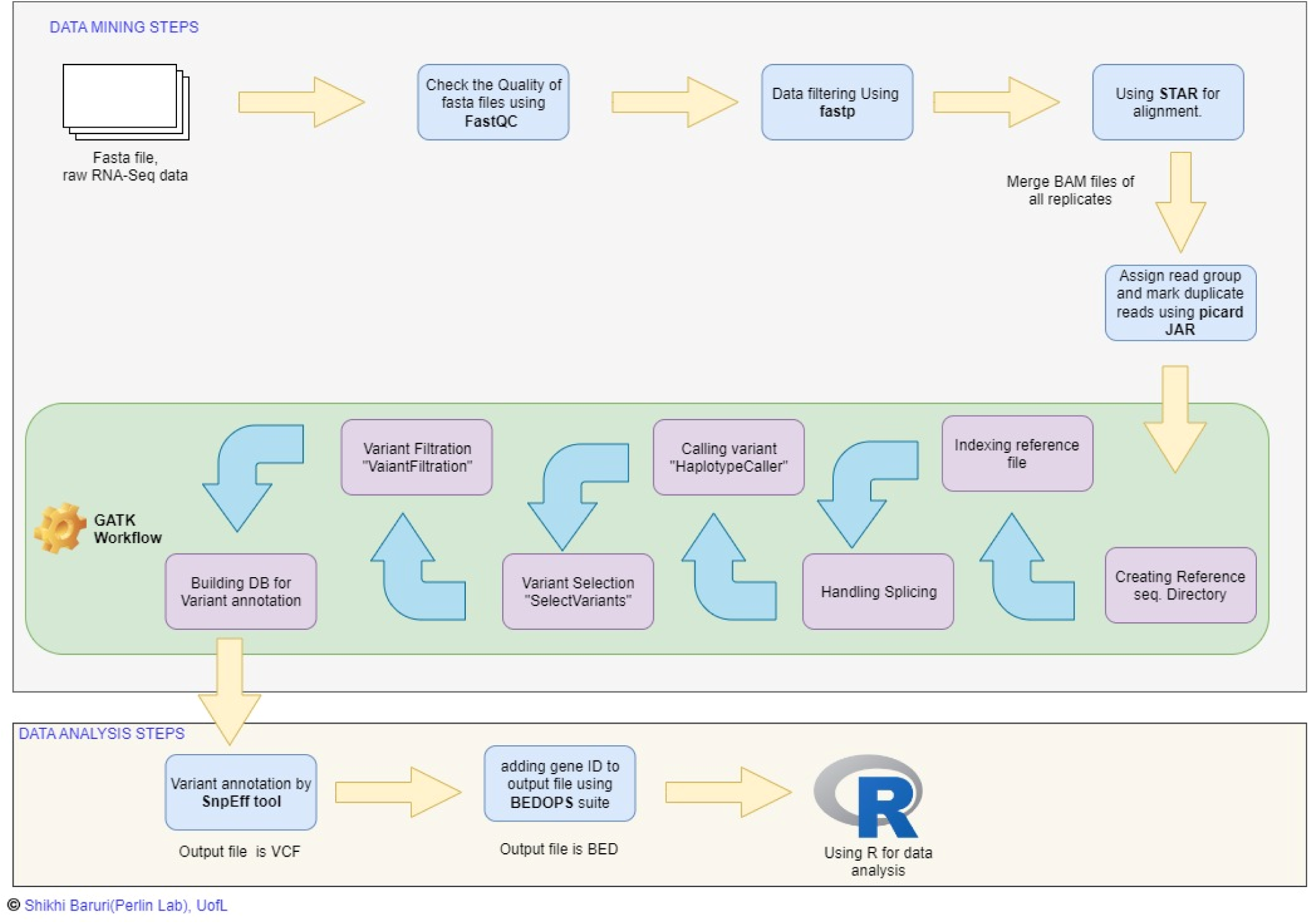
Pipeline to identify RNA editing sites.

Among the three species, MvSup, MI, and MVLG, our results suggest that the total number of edits and the proportion of edits in each haploid type differs between species. Genetic factors and environmental adaptations may influence this variability in RNA editing, highlighting the complexity of RNA editing regulation across different species. In two different mating types, a1 and a2, nonsynonymous and synonymous codon changes occurred, suggesting these codon changes may play a role in the functionality or stability of the proteins, possibly related to the necessity of environmental adaptation during a particular stage of development.

Synonymous codon changes can happen because the species have some preferred codon choice or to preserve protein function, so A to I RNA editing would therefore be necessary for these *Microbotryum* species.

When a codon change happened leading to a Thr to Ala substitution in a protein, it has a significant impact on the phosphorylation processes. Thr is a common target for phosphorylation due to a hydroxyl group present in its side chain, providing a target for addition of phosphate. In contrast, Ala lacks the hydroxyl group, so it would not be able to undergo phosphorylation [33, 34]. This alteration can affect biological processes associated with phosphorylation and signaling pathways, which regulate various cellular functions such as cell cycle progression, gene expression, and signal transduction [35].

In MvSup and MVLG, of the unique genes only edited in the haploid strains, Ala was the most common codon after nonsynonymous substitution. However, when we compare the three species, Lys to Glu conversion is the common amino acid substitution among all of the mating types. According to some studies [36], most RNA editing events result in nonsynonymous modifications; over 95% of editing events modify the amino acid sequence, which frequently makes the proteins involved more hydrophobic. Protein folding and stability can be improved by this hydrophobicity, which may be essential for the organism’s capacity to respond to environmental stressors. The reasons behind this difference could be genetic or environmental factors, which may influence the evolution or biological functions of these three species for better adaptation in the environment [36]. Furthermore, because of their sequence context and many adenosines, certain codons are more vulnerable to editing. For example, it has been discovered that the lysine codons, which feature two adenosines, are abundant in editing motifs, indicating a codon-dependent preference for editing. Because of this codon selectivity, specific amino acids, such alanine, can be preferentially expressed through A-to-I editing, which can affect the interactions and functional characteristics of the protein [37]. In the following paragraphs, several specific protein examples are discussed where A-to-I RNA editing is predicted to alter protein function during different stages of the *Microbotryum* lifecycle.

During the mating stage of MvSup, a nonsynonymous codon change occurred in a thioredoxin-like protein at codon position 18, which leads to codon changes from Thr to Ala within the functional domain of the protein. This gene is only edited during the mating stage. Thioredoxin-like protein 1 (TLP1) is an important protein in fungi to regulate cellular redox and to prevent oxidative damage [38]. Additionally, TLP1 has been detected in a number of cellular regions, including the periplasm, cytoplasm, and nucleus, suggesting its many functions in cells [39–41]. Furthermore, TLP1 is able to control redox homeostasis by its association with oxidative stress adaption in several organisms [42].

The gene associated with serine-type endopeptidase /Subtilase activity, was downregulated in the mating stage. RNA editing of that gene at the mating stage caused codon changes from Thr to Ala at position 681. In the context of subtilisin-like proteases in fungi, the substitution of a codon from Thr to Ala could potentially impact the enzymatic activity and substrate specificity of the protease [43, 44]. Subtilisin-like proteases play crucial roles in various biological processes, including plant defense, fungal virulence, and mutualistic interactions [45]. Therefore, changing the amino acid composition of these proteases in MVLG could affect the ability to cleave specific substrates, potentially affecting their biological functions. As there is a significant relationship between RNA editing and gene expression, this type of editing can significantly diversify gene expression and affect mRNA stability [46–48]. The significance of subtilisin-like proteases in fungal virulence has been emphasized, with studies reporting that these broadly distributed proteases are necessary for pathogenic fungi to be virulent [49].

A to I RNA editing occurred for the genes in common at each stage of the lifecycle of MVLG and MvSup, mostly those for transporter and transmembrane proteins. A-to-I RNA editing in RNAs of transporter proteins has yet to be extensively studied for fungi. However, transporter proteins are crucial in interactions between and among fungi and plants, particularly in mycorrhizal symbiosis [50]. Furthermore, identifying membrane proteins responsible for sensing and transporting biomass hydrolysates in anaerobic gut fungi highlights the importance of transport proteins in fungi [51]. A-to-I RNA editing in transporter proteins can be advantageous for fungi, possibly allowing them to colonize new environments and utilize previously inaccessible nutrient sources, making use of the modified forms of their transport proteins.

RNA editing of the genes only occurring in the mated stage of the three species (e.g., associated with aspartic-type endopeptidase activity/serine-type endopeptidase activity/Calcium-dependent cysteine-type endopeptidase activity), is significant. Serine proteases in symbiotic fungi play important roles in both pathogenic and mutualistic associations that emphasize their importance in fungal biology and ecology [52]. These hydrolytic enzymes are essential for the breakdown of host tissues, which makes it easier for fungi to invade and get nutrients. Serine proteases, like as subtilisin-like proteases, have been demonstrated to break down the protective plant parasitic nematodes nematode cuticle in nematophagous fungus, enabling the fungi to successfully enter and infect their hosts [53, 54].

Nonsynonymous codon changes occurred only at the mating stage in the PHB domain-containing protein that resulted in Asn to Ser and Met to Val changes, respectively. The PHB (prohibitin) domain-containing protein has been associated with cellular survival and physiology in fungi[55]. Through the Ras-Raf-MEK-ERK pathway, Prohibitin plays a significant role for cell survival. This pathway is associated with signal transduction which is vital for cell proliferation and survival. Prohibitin is thought to function as a repressor protein in this pathway, modulating the activity of downstream effectors [29].

Aspartic proteases are known for a variety of biological functions such as pathogenesis, nutrition and biocontrol. They are a group of proteolytic enzymes that function in acidic environments and share the same catalytic domain [56]. These type of enzymes are involved in various processes including tissue invasion, migration, digestion and reproduction in parasitic organisms [57]. For fungal infection they may be involved in nutritional acquisition and play an important role in breaking down the tissue barriers and in destroying the molecules generated by the host plant as a defense [58]. Also, these types of enzymes have been identified as essential components in the biocontrol activities of fungi such as *Trichoderma harzianum*, where they play an important role among the enzymes released by the fungus [59]. Moreover, with respect to pathogenesis and nutrition, aspartic proteases have been shown to be involved in the dimorphism and pathogenesis of, e.g., *Ustilago maydis*, where vacuole proteases, including aspartic proteases, have essential functions in different physiological processes in fungi [60]. Furthermore, aspartic proteases have been demonstrated to be involved in the thermo-dimorphism of *Paracoccidioides brasiliensis*, playing a crucial part in morphogenesis, cellular activity, immunology, and nutrition as well as the host invasion process [61]. In the context of plant-fungal interactions, cysteine proteases have been implicated in suppressing host immunity by inhibiting apoplastic cysteine proteases, allowing for biotrophic interactions of maize with *U. maydis* [62].

In the MVLG FIlate stage, RNA editing of a gene coding for lytic transglycoase-like enzyme degrades and remodels the fungal cell wall. These enzymes are responsible for the hydrolysis of glycosidic bonds in polysaccharides and contribute to the degradation and remodeling of fungal cell walls. These enzymes belong to the family of auxiliary activities, which catalyze the oxidative cleavage of glycosidic bonds, thereby facilitating the breakdown of complex carbohydrates into simple sugars that fungi can use [63, 64]. The presence of these enzymes in various fungal species suggests a robust mechanism for cell wall turnover, similar to the function of LTGs in bacteria. Environmental cues typically influence the production of these lytic enzymes, causing fungi to alter the composition of their cell walls and their processes of degradation. This enables the fungus to enter the plant, receiving nutrients to sustain fungal growth within the flower [24, 25, 65].

During the MIlate stage, A to I RNA editing occurred in a gene that codes for methylenomycin which is an H+ antiporter. This gene is involved in maintaining cellular ion homeostasis and moves Na+ ions out of the cytosol into the vacuole [26–28]. Furthermore, for pathogenic fungi, this methylenomycin:H+ antiporter is required for the infection process [66]. Also, this antiporter plays an important role to regulate vacuolar pH which further indicates its importance in the infection and colonization process by fungi in plants [67]. In different organisms, including plants, it is observed that this type of antiporter can catalyze the exchange of ions across the membrane [68]. Thus, RNA editing of such genes during the MIlate stage suggests a significant role in the infection process of fungi and their impact on plant health and growth.

During the FIlate stage of MVLG, A To I RNA editing occurred in a gene that codes for Peptidyl-prolyl cis-trans isomerase (PPIase)-active proteins. These types of protein are also known as cyclophilins, proteins important for protein folding, assembly, and transport. By facilitating the cis-trans isomerization of Pro residues, these proteins function as molecular chaperones and aid in the proper folding of proteins [69–73]. Within the framework of fungal-plant host-pathogen interactions, cyclophilins have been linked to the correct folding of effector proteins during infection. This folding occurs through the binding and appropriate folding of effector proteins via peptidyl-prolyl cis/trans isomerization [74]. Furthermore, by stabilizing proteins and membranes under stressful conditions, cyclophilins have been demonstrated to control plant responses to stress, including salt stress [71] [75].

In MVLG, one of the genes that was edited in all stages, codes for Ulp1 protease, a member of the ubiquitin-like protease 1 (Ulp1) cysteine protease family, which plays a crucial role in fungal mating; this gene showed more editing sites in the mated stage than in the other stages examined. In fungi, Ulp1 is essential for processing the C terminus of the Smt3 precursor into its mature form by removing its three terminal amino acids [76].

A gene that encodes an ABC type transporter was edited in all stages in both *Microbotryum* species MVLG and MvSup and has been demonstrated to play a role in drug resistance and detoxification processes in other fungi [77]. Additionally, these transporters have contributed to resistance to fungicides and mycotoxins in various fungi [77, 78]. That means this ABC type transporter may be required to adapt in each stage of both fungi for resistance.

In this study, we found that in three distinct *Microbotryum* species, nonsynonymous codon changes occurred more frequently than synonymous changes in all haploid types except one. The substitutions could be related to natural evolutionary processes such as environmental adaptations, or specific biological necessities. Overall, the study of RNA editing in Basidiomycota fungi is a relatively new and evolving field in genetics. Furthermore, the characteristics and mechanisms of RNA editing in fungi are different from those in metazoans. More research is required to fully understand the functional significance and evolutionary implications of RNA editing in fungal biology. The world of *Microbotryum* fungi, with its vast genetic and molecular landscapes, now appears to be a rich field for such scientific discovery.

## Methods

The process of identifying RNA editing sites is multi-step and primarily focuses on RNA sequencing data processing. This includes two basic steps, data mining and data analysis steps. The detailed workflow is shown in Fig 10. For the Data mining step the GATK Best Practices workflows were used which are highly regarded in the field and a gold standard [79].

### Quality Control of Raw Data

After obtaining the raw data (fastq files) from CD Genomics (Shirley, NY) or investigators, the quality of the original reads, including sequencing error rate distribution and GC content distribution, were evaluated using FastQC. The original sequencing data contain low-quality reads and adapter sequences. To ensure the quality of data analysis, raw reads were filtered to get clean reads, and the subsequent analysis was done based on clean reads. Data filtering mainly included the removal of adapter sequences in the reads, removing reads with a high proportion of N (N denotes the unascertained base information), and removing low-quality reads and other anomalies in the data. This process was carried out using fastp [80].

### Alignment to Reference Genome

In this step, the genome annotation files for MI and MVLG were downloaded from Ensemble Fungi (https://fungi.ensembl.org › index.html). For MvSup the genome annotation file was provided by R. Rodriguez de la Vega and T. Giraud.The gff file was converted into a gtf file using cufflinks packages, and then the STAR genome index was created. After the 2-pass mapping procedure, the STAR tool was used with default parameters for the read mapping on the reference genome [81].

### Post-Alignment Processing using GATK (GATK Best Practices)

After post-alignment, sorting was done using Samtools [82]. Picard (http://broadinstitute.github.io/picard/) was used to assign the read group (RG) tag and mark duplicates. The reference sequence dictionary was created, and the reference file was indexed. The SplitNCigarReads step was used to rewrite reads into a format more amenable to variant calling. This format splits reads into exon segments (getting rid of Ns) and hard-clips any sequences overhanging into the intronic regions. In the variant calling step, GATK’s HaplotypeCaller function was used for calling variants in GVCF mode to call variants from RNA-Seq data [83]. This step is vital for identifying potential RNA editing sites. After that, the "Select Variants" and "Variant Filtration" functions were used for variant selection.

In the Annotation step, we first built the database for variant annotation, and then the variants were annotated using the SnpEff tool, which provides information about each variant’s genomic context to understand the functional impact of RNA editing sites (https://gatk.broadinstitute.org/hc/en-us). Next, we converted the output vcf file into a bed file for statistical analysis.

All statistical tests were performed using R software Version 2022.12.0+353 (2022.12.0+353)[84] . REDI tool [74] was used to verify the comparison for one of the species, MI [85].

Accurate identification of A-to-I RNA editing sites in bioinformatics requires using DNA sequencing data as a control in addition to RNA sequencing data. Previous studies highlighted that to acquire an accurate profile of RNA editing sites, it is necessary to combine a transcriptome with genomic DNA resequencing of a matched sample [86]. In this study, we used DNA sequencing data as a control to identify single nucleotide polymorphisms (SNPs) sites in the genome of the strains used, compared with reference genomes. The genomic DNA was extracted from the three species of *Microbotryum*, which were grown in YPD media overnight at 28°C, followed by a PCI (phenol-chloroform-isoamyl alcohol) extraction protocol [15]. The extracted DNA was then washed with 70% ethanol. Finally, to remove any residual RNA contamination we treated the genomic DNA with PureRec RNase A enzyme (Zymo Research, Irvine, California) and further purification was done with genomic DNA clean and concentrator kit (Zymo Research) to ensure purity and quality. Then the genomic DNA of the three species were sequenced using nanopore technology (Plasmidsaurus, Louisville, Kentucky) and the SNP sites in the genomic DNA relative to reference genomes were identified. For the alignment of DNA seq data with the reference genomes the BWA-MEM2 tool was used [89–91]. The three re-sequenced genomes were submitted to NCBI and are available with the following Accession Numbers: *M. superbum* (WGS), PRJNA1201796; *M. lychnidis-dioicae* (WGS), PRJNA1202200; *M. intermedium* (WGS), PRJNA1202575. Then, the sites from RNA that matched SNPs found after DNA re-sequencing were eliminated from the inventory of RNA editing sites for each species/condition. In the case of MvSup, the RNA sequencing library was non-strand specific, and constructed with the poly(A) method. Thus, it was not feasible to determine the strand where RNA editing took place. Therefore, for this species, we consider both the A-to-G and T-to-C base changes in positive and negative-stranded genes, respectively [87].

### Statistical Analysis

In each of the three species, the proportion of RNA editing sites was analyzed between haploid types. Estimated mean proportion of RNA editing sites between a1 and a2 mating type is calculated with 95% confidence intervals. Any interval not overlapping zero means the proportion of edits in a1 differs from a2 for all of the three species. Proportional differences within each species were determined based z score and p value.

### Data availability

Whole genome shotgun sequencing, RNAseq data and genome assemblies generated for this work are available at GenBank BioProject PRJNA1169769. Accession numbers are *M. superbum* (WGS), PRJNA1201796; *M. lychnidis-dioicae* (WGS), PRJNA1202200; *M. intermedium* (WGS), PRJNA1202575. RNASeq data for all experiments listed in this section are available in GEO at NCBI, Accession Numbers/Record GSE233508 for the *M. superbum* data.

### Code availability

Relevant customized codes are available at the following link: https://github.com/shikhi18/RNAEDITING.

They use previously published packages or software, and were shared for reproducibility reasons.

### Supporting information

S1 Table. Supplementary table. Genes upregulated in the haploid stage compared to the infection stage (MIlate). (XLSX)

S2 Table. Supplementary table. Genes, identified by specific positions, uniquely edited during the mating stage compared to other stages in MvSup. (XLSX)

S3 Table. Supplementary table. Data for a single gene that underwent RNA editing at different stages and positions. (DOCX)

S4 Table. Supplementary table. Genes uniquely edited in the MIlate stage of MVLG. (XLSX)

S5 Table. Supplementary table. Genes uniquely edited in the FIlate stage of MVLG. (XLSX)

S6 Table. Supplementary table. Functions for unique genes of MIlate compared to Filate of MVLG. (XLSX)

S7 Table. Supplementary table. Genes upregulated in the haploid stage vs. MIlate of MVLG. (XLSX) S1 Figure. Supplementary figure. Differential expression level of the genes edited during MIlate stage vs haploid stage in different conditions in MVLG. (TIFF)

## Acknowledgments

We are deeply indebted to Carrie Klinge, for helpful discussions and the initial suggestion to explore RNA editing in fungi. The authors also would like to thank Ricardo Rodriguez de la Vega and Tatiana Giraud for supplying genomic sequence information for MvSup and Roxanne Hayes for providing raw RNA seq data for MvSup and MI. We are also grateful to two anonymous reviewers whose constructive criticisms led to an improved final paper. This work was supported by National Science Foundation (NSF) Award# 2007449, NSF Award# 2419465, and USDA National Institute of Food and Agriculture (NIFA) Award#2024-67014-43676 to MHP.

## Author Contributions

**Conceptualization:** Shikhi Baruri, Michael H. Perlin

**Data curation:** Shikhi Baruri, Joseph P. Ham.

**Formal analysis:** Shikhi Baruri, Alycia Lackey.

**Funding acquisition:** Michael H. Perlin.

**Investigation:** Shikhi Baruri, Joseph P. Ham.

**Methodology:** Shikhi Baruri, Joseph P. Ham

**Project administration:** Michael H. Perlin.

**Resources:** Shikhi Baruri, Joseph P. Ham, Alycia Lackey.

**Supervision:** Michael H. Perlin

**Validation:** Shikhi Baruri, Joseph P. Ham, Alycia Lackey.

**Visualization:** Shikhi Baruri.

**Writing – original draft:** Shikhi Baruri.

**Writing – review & editing:** Shikhi Baruri, Alycia Lackey, Michael H. Perlin.

## Notes

### Competing Interest Statement

The authors have declared no competing interest.

